# Rapid *in vitro* prototyping of O-methyltransferases for pathway applications in *Escherichia coli*

**DOI:** 10.1101/2020.08.27.258715

**Authors:** Kristina Haslinger, Thomas Hackl, Kristala L.J. Prather

## Abstract

O-methyltransferases are ubiquitous enzymes involved in biosynthetic pathways for secondary metabolites such as bacterial antibiotics, human catecholamine neurotransmitters, and plant phenylpropanoids. While thousands of putative O-methyltransferases are found in sequence databases, few examples are functionally characterized. From a pathway engineering perspective, however, it is crucial to know the substrate and product ranges of the respective enzymes to fully exploit their catalytic power.

In this study, we developed an *in vitro* prototyping workflow that allowed us to screen ~30 enzymes against five substrates in three days with high reproducibility. We combined *in vitro* transcription/translation of the genes of interest with a microliter-scale enzymatic assay in 96-well plates. The substrate conversion was indirectly measured by quantifying the consumption of the S-adenosyl-L-methionine co-factor by time-resolved fluorescence resonance energy transfer rather than time-consuming product analysis by chromatography. This workflow allowed us to rapidly prototype thus-far uncharacterized O-methyltransferases for future use as biocatalysts.

## Introduction

Methylation of secondary metabolites is a prevalent reaction that alters the bioavailability and reactivity of molecules^1^. This effect is important for the native function of secondary metabolites for the producer organism but also for pharmaceutical and nutraceutical applications of natural products. One example is the oxygen-directed methylation (O-methylation) of the lignin precursor caffeic acid towards ferulic acid. This reaction is crucial for regulating the rigidity of lignified cell walls in vascular plants^2^ and has been described to modulate the cytotoxicity and radical scavenging properties of isolated phenolic acids when tested for pharmaceutical applications such as neuroprotection^3,4^. Similar observations were made for methylated flavonoids (plants)^5,6^, antimicrobial peptides (bacteria)^7,8^ and dopamine (humans)^9^. O-methylation in nature is carried out by methyltransferases under the utilization of S-adenosyl-L-methionine (SAM) as an electron-deficient methyl donor thereby forming S-adenosyl-L-homocysteine. Some O-methyltransferase (OMT) families additionally require the presence of metal ions such as Mg^2+^. For OMTs acting on small molecules (excluding nucleic acids and proteins), there are several protein families with distinct sequence motifs and with a remarkable breadth in functionality. The functional exploration of these families has been somewhat anecdotal to date and has been very much focused on plant enzymes of the methyltransferase families 2 (PF00891) and 3 (PF01596)^1^. This can most likely be attributed to the fact that already in the pre-genomic era, these plant enzymes had been studied with biochemical methods^10,11^. However, with the rapid expansion of genomes sequenced to date, the methyltransferase protein families are growing by the minute and functional studies are lagging behind^12^.

In the last decade rapid advances in parallelization of molecular cloning, enzymatic assays, and even fermentation through liquid handling technologies and automation have greatly increased the throughput of functional studies of enzyme libraries^13,14^. However, the bottlenecks in these screening pipelines remain the failing of molecular cloning steps and the throughput of the reaction readout in the absence of colorimetric or fluorometric assays, which requires time-consuming chromatography methods to analyze the products^15,16^. To overcome these hurdles in the functional screening of SAM-dependent methyltransferases, we developed a rapid *in vitro* prototyping workflow to express and functionally screen a range of O-methyltransferases against several substrates. To minimize time and effort spent on molecular cloning, we employed a recently developed *in vitro* transcription/translation platform for linear DNA templates with high enzyme yields (myTXTL^®^)^17,18^, and combined it with a fluorescence-based read-out^19,20^ to monitor the consumption of the SAM co-factor. For one substrate we translated the newly gained knowledge into the development of a microbial cell factory to produce ferulic acid from simple building blocks.

## Results

### Design and benchmarking of the prototyping workflow

In order to facilitate the fast screening of a library of putative O-methyltransferases against several substrates, we set out to develop a prototyping method that is rapid and parallelizable. Herein, we identified the detection of enzymatic activity and the cloning and expression of the genes of interest as the two major bottlenecks. For the detection of enzymatic activity, we deemed a desirable approach to be independent of the substrates and products and to not require time-consuming chromatography. We turned towards commercially available assays to detect the consumption of the SAM cofactor and decided to use the TR-FRET Bridge-It^®^ S-Adenosyl Methionine (SAM) Fluorescence Assay Kit from Mediomics LLC (St. Louis, Missouri). In this endpoint assay, SAM binds to a DNA-binding protein and induces the association of two fluorescently labeled DNA fragments (donor and acceptor) for Time-Resolved Foerster Resonance Energy Transfer (TR-FRET) to occur. We hypothesized that in this way, several enzymes could be screened against multiple substrates in parallel in the plate reader, with lower TR-FRET readings observed for active enzyme-substrate combinations. We set out to test this detection method with recombinantly expressed and purified MxSafC, an enzyme known to catalyze the methylation of caffeic acid^25,32^, and compare the TR-FRET read-out with product analysis by HPLC. Compared to the control reaction without enzyme, we saw consumption of caffeic acid and SAM, as measured by HPLC and TR-FRET, respectively, after 3h of incubation (Figure 1a). We observed a good correlation of the biological replicates within and across both detection methods with a slight over-estimation of substrate-consumption with the TR-FRET assay compared to HPLC detection.

**Figure 1:**
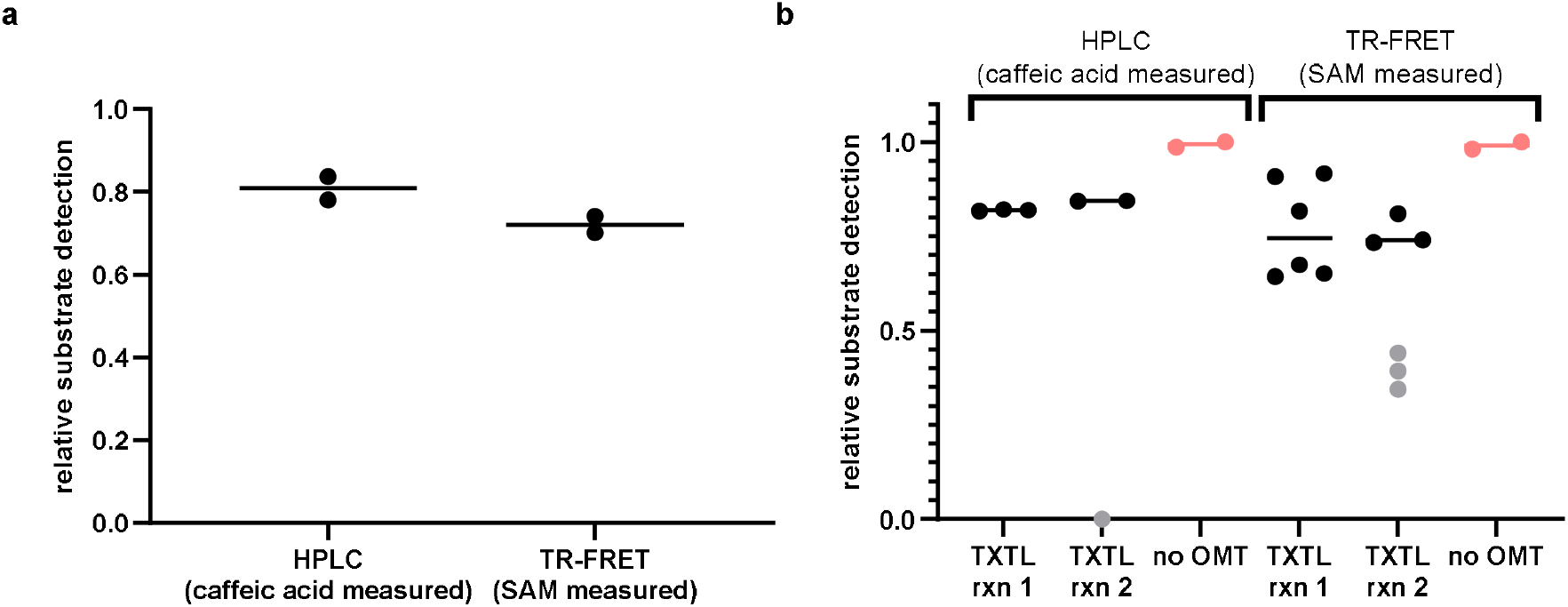
Substrate detection at the end of the OMT reaction with MxSafC as a catalyst relative to the negative controls measured by HPLC (caffeic acid concentration measured) and TR-FRET (SAM concentration measured). a) OMT reaction performed with recombinantly expressed and purified MxSafC (data points are biological replicates (n=2, median)). b) OMT reaction performed with MxSafC expressed from linear DNA in the myTXTL *in vitro* transcription/translation kit (two biological replicates are shown in separate columns; data points within each column are technical replicates of the OMT reaction and the TR-FRET assay; “no OMT” data points are biological replicates; outliers (grey data points) are likely caused by a pipetting error in the OMT assay and were excluded from determining the median).

In order to address the cloning and expression bottleneck, we decided to use an *in vitro* transcription/translation expression platform, myTXTL^®^ from Arbor Bioscience (Ann Arbor, Michigan). This allowed us to express the genes of interest from synthetic, linear DNA fragments and saved us additional time for cloning, sequence verification, transformation, protein expression and cell lysis (~up to 3 weeks of work). The linear DNA fragments were designed to contain a σ70 promoter, a T500 terminator and flanking overhangs of about 500bp to protect from degradation in the myTXTL^®^ mix. Additionally, GamS protein was added to the reactions to protect the DNA fragments. We first tested the compatibility of the myTXTL^®^ reaction mix with the OMT assay and the TR-FRET detection method with MxSafC expressed from a linear template (Figure 1b). We performed two TXTL reactions at 29°C over night and split them into three OMT reactions each. After stopping the OMT reactions, we analyzed them with the TR-FRET assay (two technical replicates) and HPLC. As negative controls we included two TXTL reactions that did not contain OMT-encoding DNA template (no technical replicates). Looking at the median of the data points, we again observe good correlation of the replicates within and cross the detection methods however, the TR-FRET assay appears to be more sensitive to experimental error than the HPLC detection. The biological replicates of the enzyme expression (TXTL reactions 1 and 2) show only minor deviation, indicating that the experimental error in the expression step of the workflow is minimal.

Since we observed good correlation between the two detection methods, yet low overall turnover yields, we proceeded with the established workflow with an extended incubation time for the OMT reaction (24h) in later experiments.

### Screening of putative OMTs for methylation of caffeic acid

In order to diversify our knowledge of OMTs in organisms other than plants, the premise of this study was to characterize a range of putative OMTs from various non-plant donor organisms across a relatively wide sequence landscape. Therefore, we first identified distantly related OMTs in the NCBI reference proteome database based on Hidden Markov Models (HMM) constructed from known plant caffeic acid OMTs^23,24^ (search input 1, Extended Data Table 1), and bacterial OMTs previously found to have a broad substrate tolerance towards catechols^25–28^ (search input 2). We found 15,994 unique sequences from all kingdoms of life, from ~190 PFAM families (Supplementary File 1). About 85% of the sequences were annotated as methyltransferases, more specifically 82% as OMTs, and about 10% contained dimerization domains. We filtered the sequences by length and alignment score and constructed a sequence similarity network to group them into clusters by pairwise amino acid sequence similarity. From the thus generated clusters, putative OMTs were chosen for experimental characterization (Table 1, Extended Data Figure 1) by taking the following criteria into consideration: the ranking of HMM scores within the clusters, a wide taxonomic spread over the selected enzymes and a balanced selection of enzymes found with the two HMMsearch runs. Multiple enzymes were chosen from the major clusters but also some high scoring putative OMTs were picked from the smallest clusters.

**Table 1:**
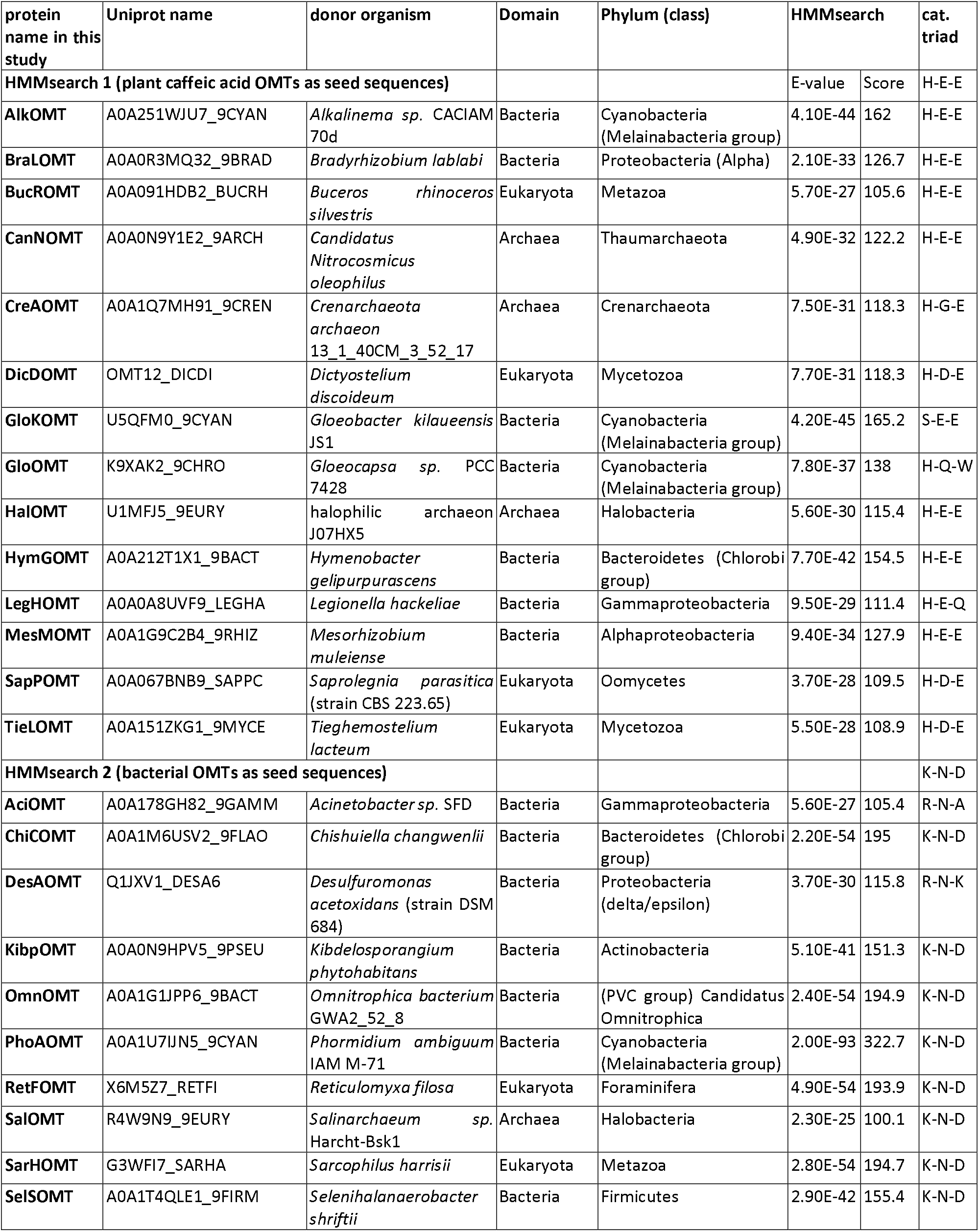
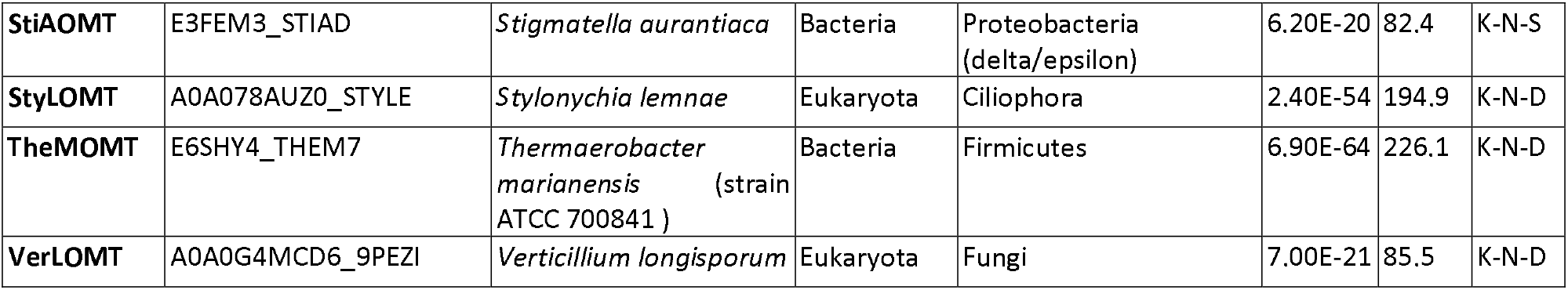
List of putative OMT selected for experimental characterization.

| protein name in this study | Uniprot name | donor organism | Domain | Phylum (class) | HMMsearch |  | cat. triad |
| --- | --- | --- | --- | --- | --- | --- | --- |
| HMMsearch 1 (plant caffeic acid OMTs as seed sequences) |  |  |  |  | E-value | Score | H-E-E |
| AlkOMT | A0A251WJU7_9CYAN | *Alkalinema* sp. CACIAM 70d | Bacteria | Cyanobacteria (Melainabacteria group) | 4.10E-44 | 162 | H-E-E |
| BraLOMT | A0A0R3MQ32_9BRAD | *Bradyrhizobium lablabi* | Bacteria | Proteobacteria (Alpha) | 2.10E-33 | 126.7 | H-E-E |
| BucROMT | A0A091HDB2_BUCRH | *Buceros rhinoceros silvestris* | Eukaryota | Metazoa | 5.70E-27 | 105.6 | H-E-E |
| CanNOMT | A0A0N9Y1E2_9ARCH | *Candidatus Nitrocosmicus oleophilus* | Archaea | Thaumarchaeota | 4.90E-32 | 122.2 | H-E-E |
| CreAOMT | A0A1Q7MH91_9CREN | *Crenarchaeota archaeon* 13_1_40CM_3_52_17 | Archaea | Crenarchaeota | 7.50E-31 | 118.3 | H-G-E |
| DicDOMT | OMT12_DICDI | *Dictyostelium discoideum* | Eukaryota | Mycetozoa | 7.70E-31 | 118.3 | H-D-E |
| GloKOMT | U5QFM0_9CYAN | *Gloeobacter kilaeensis* JS1 | Bacteria | Cyanobacteria (Melainabacteria group) | 4.20E-45 | 165.2 | S-E-E |
| GloOMT | K9XAK2_9CHRO | *Gloeocapsa* sp. PCC 7428 | Bacteria | Cyanobacteria (Melainabacteria group) | 7.80E-37 | 138 | H-Q-W |
| HalOMT | U1MFJ5_9EURY | halophilic archaeon J07HX5 | Archaea | Halobacteria | 5.60E-30 | 115.4 | H-E-E |
| HymGOMT | A0A212T1X1_9BACT | *Hymenobacter gelipurpurascens* | Bacteria | Bacteroidetes (Chlorobi group) | 7.70E-42 | 154.5 | H-E-E |
| LegHOMT | A0A0A8UVF9_LEGHA | *Legionella hackeliae* | Bacteria | Gammaproteobacteria | 9.50E-29 | 111.4 | H-E-Q |
| MesMOMT | A0A1G9C2B4_9RHIZ | *Mesorhizobium muleiense* | Bacteria | Alphaproteobacteria | 9.40E-34 | 127.9 | H-E-E |
| SapPOMT | A0A067BNB9_SAPPC | *Saprolegnia parasitica* (strain CBS 223.65) | Eukaryota | Oomycetes | 3.70E-28 | 109.5 | H-D-E |
| TieLOMT | A0A151ZKG1_9MYCE | *Tieghemostelium lacteum* | Eukaryota | Mycetozoa | 5.50E-28 | 108.9 | H-D-E |
| HMMsearch 2 (bacterial OMTs as seed sequences) |  |  |  |  |  |  | K-N-D |
| AciOMT | A0A178GH82_9GAMM | *Acinetobacter* sp. SFD | Bacteria | Gammaproteobacteria | 5.60E-27 | 105.4 | R-N-A |
| ChiCOMT | A0A1M6USV2_9FLAO | *Chishuiella changwenlii* | Bacteria | Bacteroidetes (Chlorobi group) | 2.20E-54 | 195 | K-N-D |
| DesAOMT | Q1JXV1_DESA6 | *Desulfuromonas acetoxidans* (strain DSM 684) | Bacteria | Proteobacteria (delta/epsilon) | 3.70E-30 | 115.8 | R-N-K |
| KibpOMT | A0A0N9HPV5_9PSEU | *Kibdelosporangium phytohabitans* | Bacteria | Actinobacteria | 5.10E-41 | 151.3 | K-N-D |
| OmnOMT | A0A1G1JPP6_9BACT | *Omnitrophica bacterium* GWA2_52_8 | Bacteria | (PVC group) Candidatus Omnitrophica | 2.40E-54 | 194.9 | K-N-D |
| PhoAOMT | A0A1U7IJN5_9CYAN | *Phormidium ambiguum* IAM M-71 | Bacteria | Cyanobacteria (Melainabacteria group) | 2.00E-93 | 322.7 | K-N-D |
| RetFOMT | X6M5Z7_RETFI | *Reticulomyxa filosa* | Eukaryota | Foraminifera | 4.90E-54 | 193.9 | K-N-D |
| SalOMT | R4W9N9_9EURY | *Salinarchaeum* sp. Harcht-Bsk1 | Archaea | Halobacteria | 2.30E-25 | 100.1 | K-N-D |
| SarHOMT | G3WFI7_SARHA | *Sarcophilus harrisii* | Eukaryota | Metazoa | 2.80E-54 | 194.7 | K-N-D |
| SelSOMT | A0A1T4QLE1_9FIRM | *Selenihalanaerobacter shriftii* | Bacteria | Firmicutes | 2.90E-42 | 155.4 | K-N-D |
| **StiAOMT** | E3FEM3_STIAD | *Stigmatella aurantiaca* | Bacteria | Proteobacteria<br>(delta/epsilon) | 6.20E-20 | 82.4 | K-N-S |
| **StyLOMT** | A0A078AUZ0_STYLE | *Stylonychia lemnae* | Eukaryota | Ciliophora | 2.40E-54 | 194.9 | K-N-D |
| **TheMOMT** | E6SHY4_THEM7 | *Thermaerobacter marianensis* (strain ATCC 700841 ) | Bacteria | Firmicutes | 6.90E-64 | 226.1 | K-N-D |
| **VerLOMT** | A0A0G4MCD6_9PEZI | *Verticillium longisporum* | Eukaryota | Fungi | 7.00E-21 | 85.5 | K-N-D |

The first large-scale screen of the selected putative OMTs was performed with caffeic acid as the substrate. We analyzed the enzymatic reactions by HPLC and TR-FRET (Figure 2a,b) and repeated the experiment on a different day with a slightly different sample-handling workflow that allowed the consistent use of multichannel pipettes throughout the experiment (Figure 2c,d). The data points in the plots are ordered by increasing substrate turnover based on panel C. The HPLC analysis shows a good correlation of the independent experiments with each other, both in relative terms (ranking of the tested enzyme by performance) and in absolute terms. This again indicates that the expression levels in the TXTL reactions is highly reproducible and that the technical error in the OMT reaction is low. However, for the TR-FRET analysis of the first experiment (Figure 2b), it is evident that the technical error by manual sample dilution and setup of the TR-FRET detection assay is very high and therefore, the technical replicates deviate strongly. The overall noise of the experiment is very high, which becomes most apparent in the wells that appear to have higher SAM concentrations than the negative controls (here shown at the bottom of the plot). These experimental errors were overcome with a slightly different sample handling procedure in the second experiment (Figure 2d), which shows dramatically decreased noise in the data and a clear distinction between true- and false-positives. However, even in the first experiment with high background noise the best-performing enzymes can be clearly distinguished from the other ones. In the intermediate range, it is difficult to make a distinct cut-off. However, depending on the goal of this screening step, the cut-off can be set at a lower or higher level of SAM consumption at the risk of including false-positives or excluding false-negatives, respectively. In this case, we decided to make a very conservative cut-off and to even carry some true-negatives forward to the characterization in *E. coli* (*vide infra*). Overall, we observe a clear correlation between the HPLC an TR-FRET read-out and were therefore encouraged to screen the enzymes against four other potential substrates: 1,2-dihydroxybenzene (catechol), ferulic acid, quercetin and dopamine.

**Figure 2:**
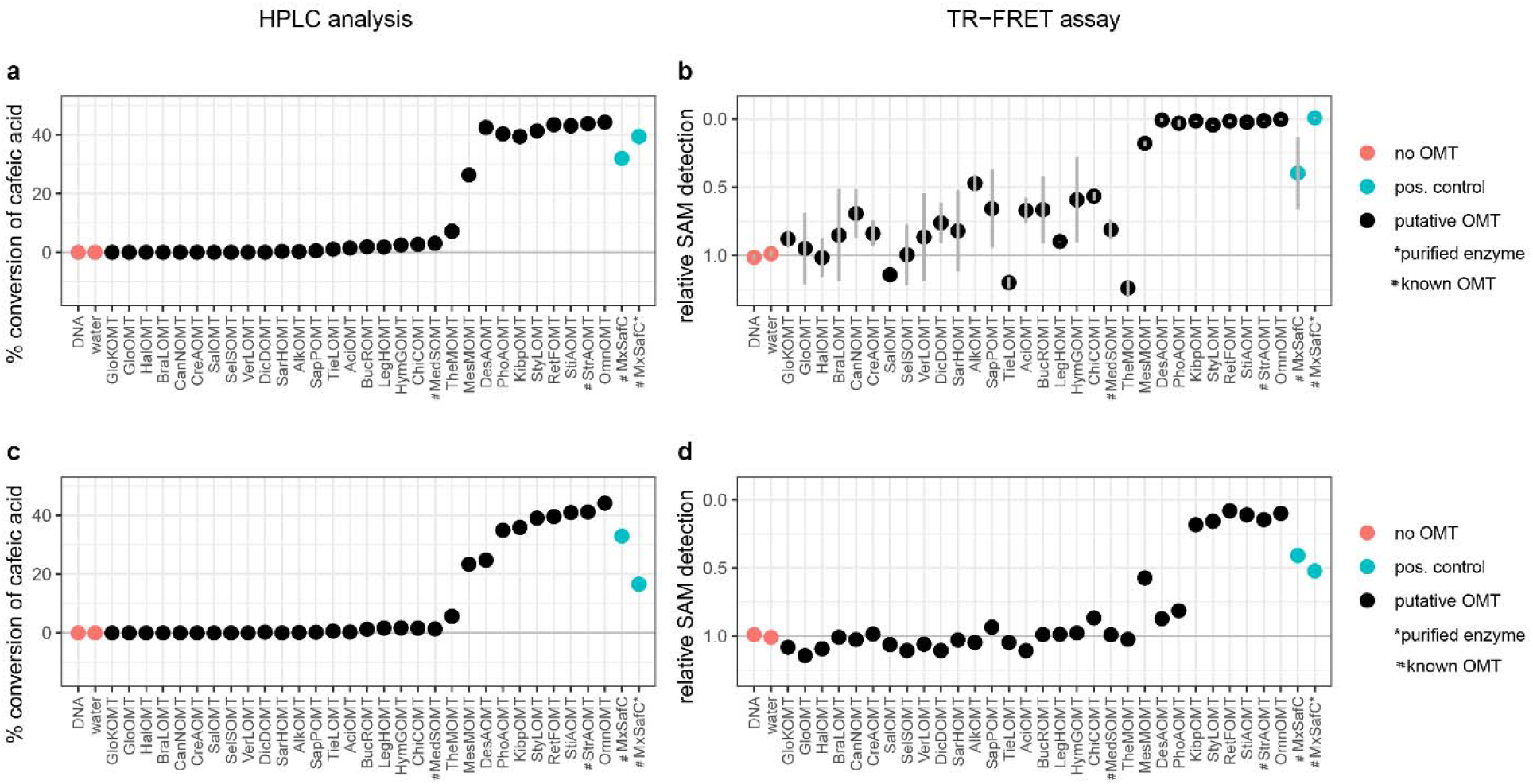
Screening of 30 enzymes of interest against caffeic acid as substrate and SAM as co-substrate in two independent experiments. OMT reaction stopped after 24h. a) and c) Samples analyzed by HPLC, expressed as percent caffeic acid converted. b) and d) Samples analyzed by TR-FRET expressed as SAM levels detected relative to the “no OMT” controls; b) technical replicates of the TR-FRET assay shown as mean +/− SD, n=2; d) single measurement. Data points sorted by increasing substrate turnover based on panel c; Red data points – negative controls, blue data points – positive controls.

### Screening of putative OMTs against other substrates

Next, we sought to use our *in vitro* expression and testing workflow to screen our panel of putative OMTs against other substrates. We selected catechol and dopamine - two known substrates for MxSafC and plant caffeic acid OMTs-, quercetin – a flavonoid also often converted by plant caffeic acid OMTs -, and ferulic acid – the precursor for a non-natural double-methylated product. We ran all reactions in parallel by diluting the TXTL reactions after overnight expression and aliquoting them into microtiter plates with the OMT reaction mixes. After 24h we stopped the OMT reactions and assessed the SAM levels with the TR-FRET assay. We observed increased SAM consumption by 11 OMTs in the presence of catechol (Figure 3a) and by 15 OMTs in the presence of dopamine (Figure 3b), whereas in the presence of ferulic acid and quercetin, only low levels of SAM conversion were observed that are difficult to separate from the background noise of the assay (Figure 3c and d). Furthermore, we did not have any true positive controls for these substrates in the panel of enzymes. Therefore, we are inclined to interpret the results as negative for these substrates. Also, in the presence of dopamine, the separation of positives and negatives is less clear-cut than with catechol. However, since the background noise appears to be rather small, we’d suggest a more inclusive cut-off for further analysis.

**Figure 3:**
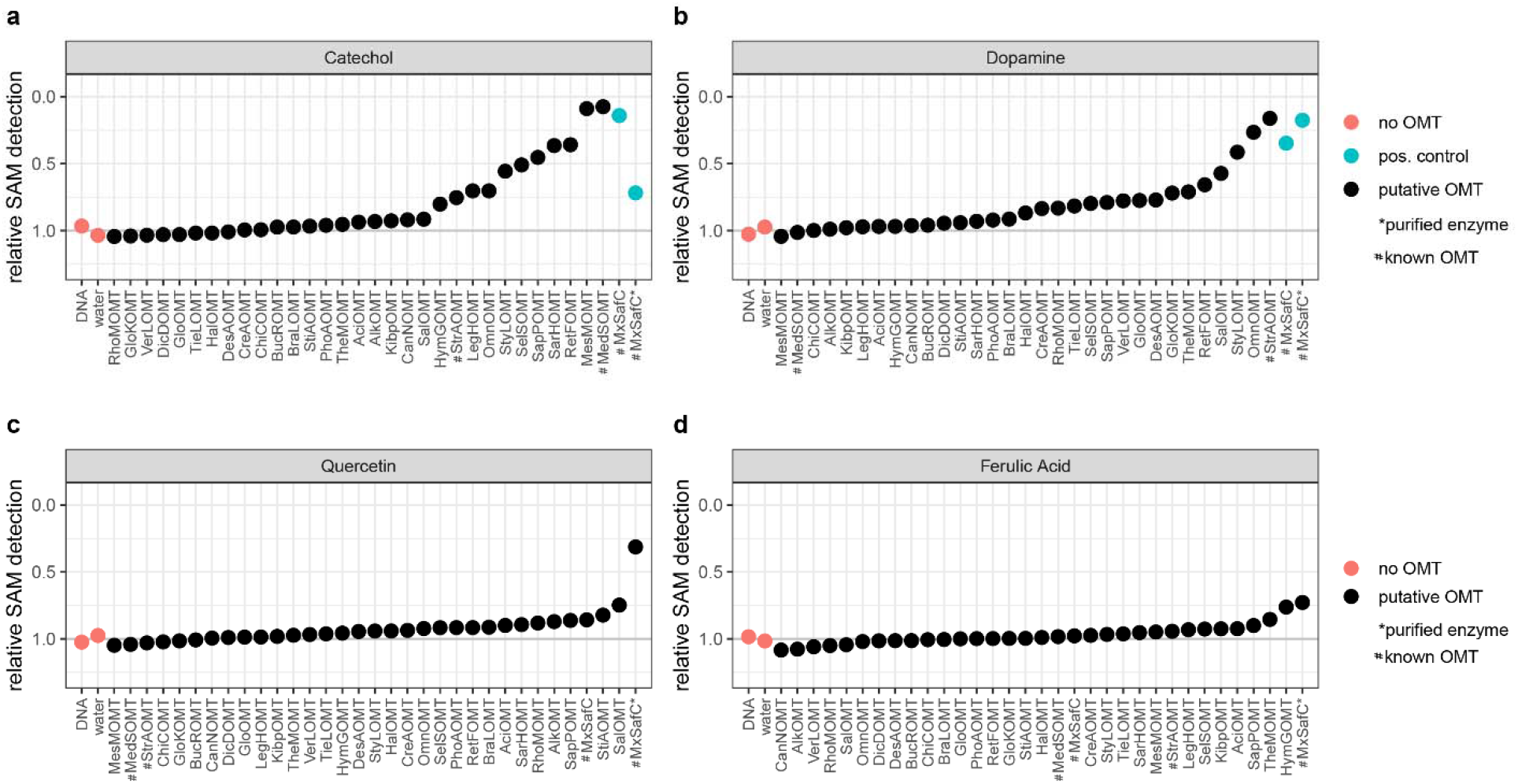
Screening for OMT activity of all enzymes of interest *in vitro* with four different substrates. a) catechol, b) dopamine, c) quercetin and d) ferulic acid. Data points ordered by increasing substrate conversion for each panel from left to right; Red data points – negative controls, blue data points – positive controls (if available).

Only considering the enzymes with highest SAM conversion, we see overlap in substrate acceptance for StyLOMT, RetFOMT, StrAOMT, OmnOMT and MesMOMT. While the former enzymes show increased SAM conversion in the presence of all three substrates, MesMOMT is not stimulated by dopamine. Several enzymes appear to display stronger substrate selectivity: KibPOMT and StiAOMT are selective for caffeic acid, HymGOMT, LegHOMT, MedSOMT, SapPOMT, SarHOMT and SelSOMT are selective for catechol, and SalOMT is selective for dopamine. A sequence comparison of the tested OMTs shows that enzymes with similar activities also share higher sequence similarity with each other (Figure 4). Looking at the active site residues predicted based on multiple sequence alignments, we see that some of the tested OMTs display activity although they carry changes in the putative catalytic triad (Table 1). In the group of enzymes from HMMsearch 1 (plant input sequences) the catalytic triad should be H-E-E (in *Medicago sativa* COMT1_MEDSA residues H269, E297 and E329^39^) and is highly conserved with a few exceptions: CreAOMT, GloKOMT, GloOMT, LegHOMT, with all enzymes being active. In the group of enzymes from HMMsearch 2 (bacterial input sequences) the catalytic triad should be K-N-D (in MxSafC residues K145, N69, D212^40^) and is even more conserved. Only three sequences AciOMT, DesAOMT and StiAOMT show significant changes in these amino acids, with only AciOMT being inactive. This suggests that there must be other changes in the active site architecture of these OMTs that compensate for these amino acid substitutions. With a stricter pre-selection based on sequence similarity and active site conservation, we might have missed these interesting OMTs, whereas our pre-screening approach enabled us to explore a wider sequence space.

**Figure 4:**
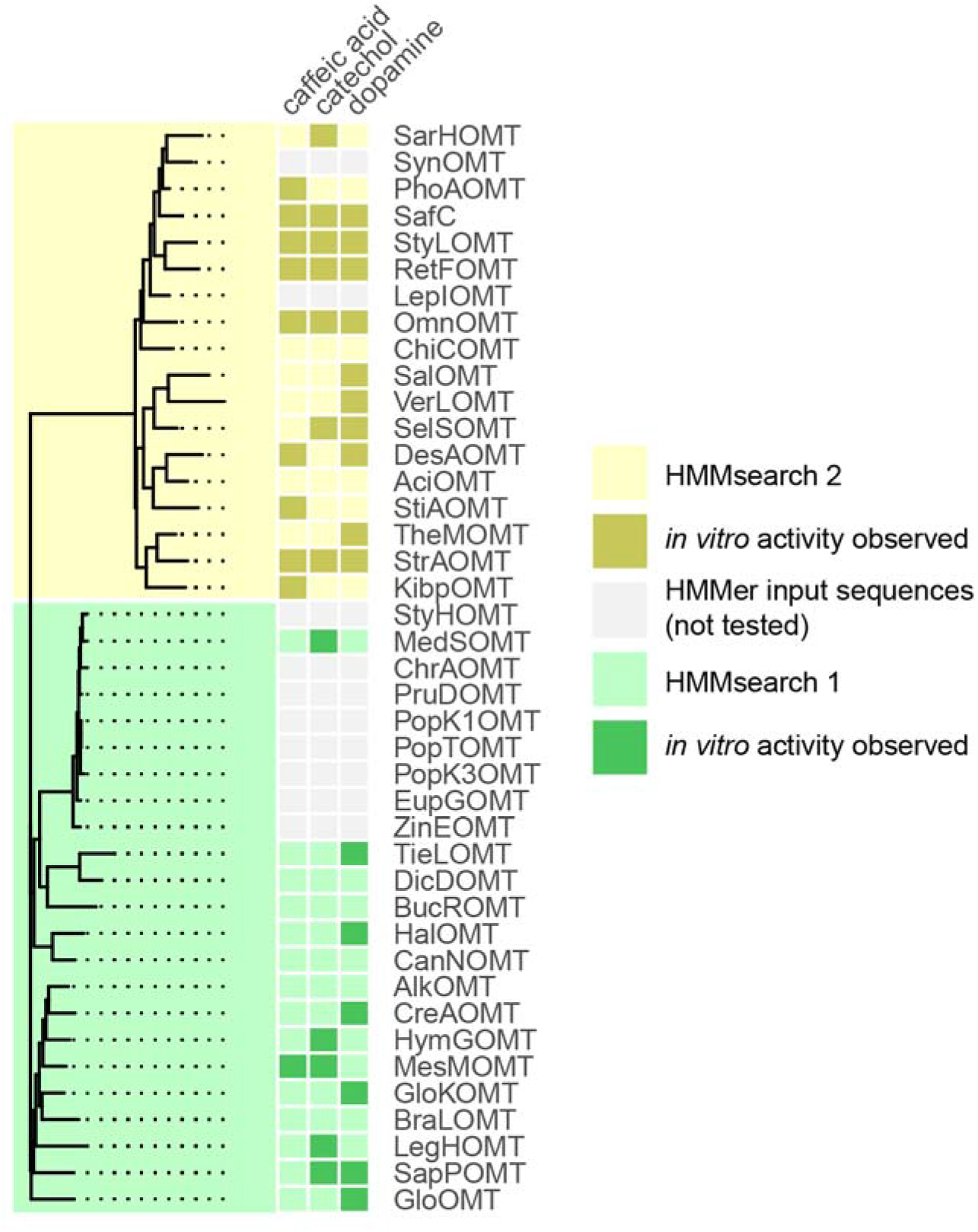
Phylogenetic tree based on a multiple sequence alignment of the enzymes screened in this study and the HMMsearch input sequences (left; green: HMMsearch1; yellow: HMMsearch 2) side-by-side with the *in vitro* activity data on caffeic acid, catechol and dopamine (right, dark color=*in vitro* activity observed).

### Application of pre-screened OMTs in a pathway towards (iso-)ferulic acid

Lastly, we sought to use the pre-screened OMTs in an *E. coli* microbial cell factory. We chose to expand our previously constructed and optimized pathway from tyrosine to caffeic acid^31,41^ by one enzymatic step in order to generate the pharmaceutically relevant phenolic acid, ferulic acid and it’s regio-isomer 4-methoxy-3-hydroxy-cinnamic acid (iso-ferulic acid). In order to select enzymes for testing in the recombinant pathway, we first explored the data from the pre-screening assay in the context of enzyme expression, protein sequence and the donor organism. We found that the presence or absence of a band of the appropriate size in the SDS PAGE did not correlate with observed enzymatic activity (Extended Data Figure 2). For instance, in the lane of one of the best performing enzyme, StyLOMT, we did not see a distinct band on SDS PAGE. Whereas for some inactive enzymes such as, HalOMT, we saw a distinct band on SDS PAGE. This indicates that some enzymes are expressed at a low level yet active, whereas others are either not correctly expressed and folded, or were simply not challenged with the right substrate in this study. When mapping the pre-screening results onto the sequence similarity network we notice that the enzymes active on caffeic acid (Extended Data Figure 1, filled symbol) are distributed across the network with most of them being part of the main cluster. All active OMTs except MesMOMT show highest sequence similarity with the bacterial seed sequences (yellow box). This indicates that the bacterial input sequences provided a better search template for identifying new caffeic acid OMTs than the plant input sequences. Since some of the putative OMTs found with the plant input sequences display activity against catechol and dopamine, we can exclude that the lack of activity on caffeic acid is caused by a general problem with our *in silico* selection, *in vitro* expression and pre-screening approach. However, for the putative OMTs that did not display activity on any of the tested substrates, we cannot rule out protein expression and folding problems. Two of the active enzymes, RetFOMT and StyLOMT, are from eukaryotic donors and the rest from bacterial donors. This indicates that the pre-screening method is also applicable to eukaryotic enzymes.

Based on the prescreening results, we chose the ten top-performing enzymes including the previously characterized StrAOMT^28^ and MxSafC^25,32^ and two enzymes that were inactive in the pre-screen: HalOMT (archaeal donor, visibly expressed) and SalOMT (archaeal donor, not visibly expressed). We cloned the respective genes into the vector pRSFduet::FjTAL, which already contained the tyrosine ammonia lyase gene from *Flavobacterium johnsoniae* (FjTAL, first pathway step) in a separate expression cassette, for expression under the T7 promoter (Extended Data Table 2). We co-transformed each new plasmid with two other plasmids encoding for the Cytochrome P450 monooxygenase CYP199A2 F185L NΔ7 and its redox partners (second pathway step) from our previous study^31^ into *E. coli* K12 MG1655DE3. In the resulting strains (s01-s12, Extended Data Table 3) L-tyrosine will be converted to *p*-coumaric acid by FjTAL, to caffeic acid by CYP199A2 F185L NΔ7 and to (iso-)ferulic acid by the OMTs. As a negative control, we used a strain with the pRSF::FjTAL plasmid lacking an OMT gene (s00). In initial fermentation experiments with the modified M9 minimal media composition that we had previously used^31^, we did not observe significant product formation from glucose or fed L-tyrosine (data not shown) and therefore decided to first optimize the conditions for the OMT catalyzed step with a subset of the strains and in smaller scale reactions with fed caffeic acid. We observed that the addition of Mg^2+^ (obligate co-factor for OMTs) by itself only led to slightly higher caffeic acid conversion, whereas feeding of L-methionine as a precursor for SAM improved the turnover by 2.2- to 3.6-fold (Figure 5a). This finding is consistent with previous observations for vanillin biosynthesis in *E. coli*^42^ and indicates that SAM supply is limited and needs to be increased for OMT containing pathways to be efficient. With this knowledge, we tested all strains in 15 mL fermentations with glucose as a carbon source, L-tyrosine as a pathway precursor and Mg^2+^ and L-methionine as additives for the OMT reaction. We observed product formation for all strains expressing OMTs that had tested active in the pre-screening step (Figure 5b). In most strains more than half of the caffeic acid formed was converted to the methylated products and four strains even achieved full conversion: s06 expressing PhoAOMT, s08 expressing StyLOMT, s09 expressing RetFOMT and s11 expressing StrAOMT. In terms of titers, s11 displays the most desirable outcome with low titers for pathway intermediates and side products, and a high product titer of 0.49 mM +/− 0.06 mM (Figure 5c). Interestingly, all OMTs displayed a strong regioselectivity for the meta-position over the para-position *in vivo*, although some showed a preference for the para-position in the *in vitro* screening step (Extended Data Table 4). This indicates that *in vitro* data cannot necessarily be directly translated into whole-cell applications. Nevertheless, our pre-screening step decreased the experimental load for cloning, fermentation and product analysis by HPLC by at least two-thirds. The best performing OMT, StrAOMT, had previously been observed to act on caffeic acid, however with low catalytic efficiency^28^. To our surprise it was one of the top-performers in the pre-screening and the pathway application in this study. To the best of our knowledge, StrAOMT has not been used in the context of a pathway before.

**Figure 5:**
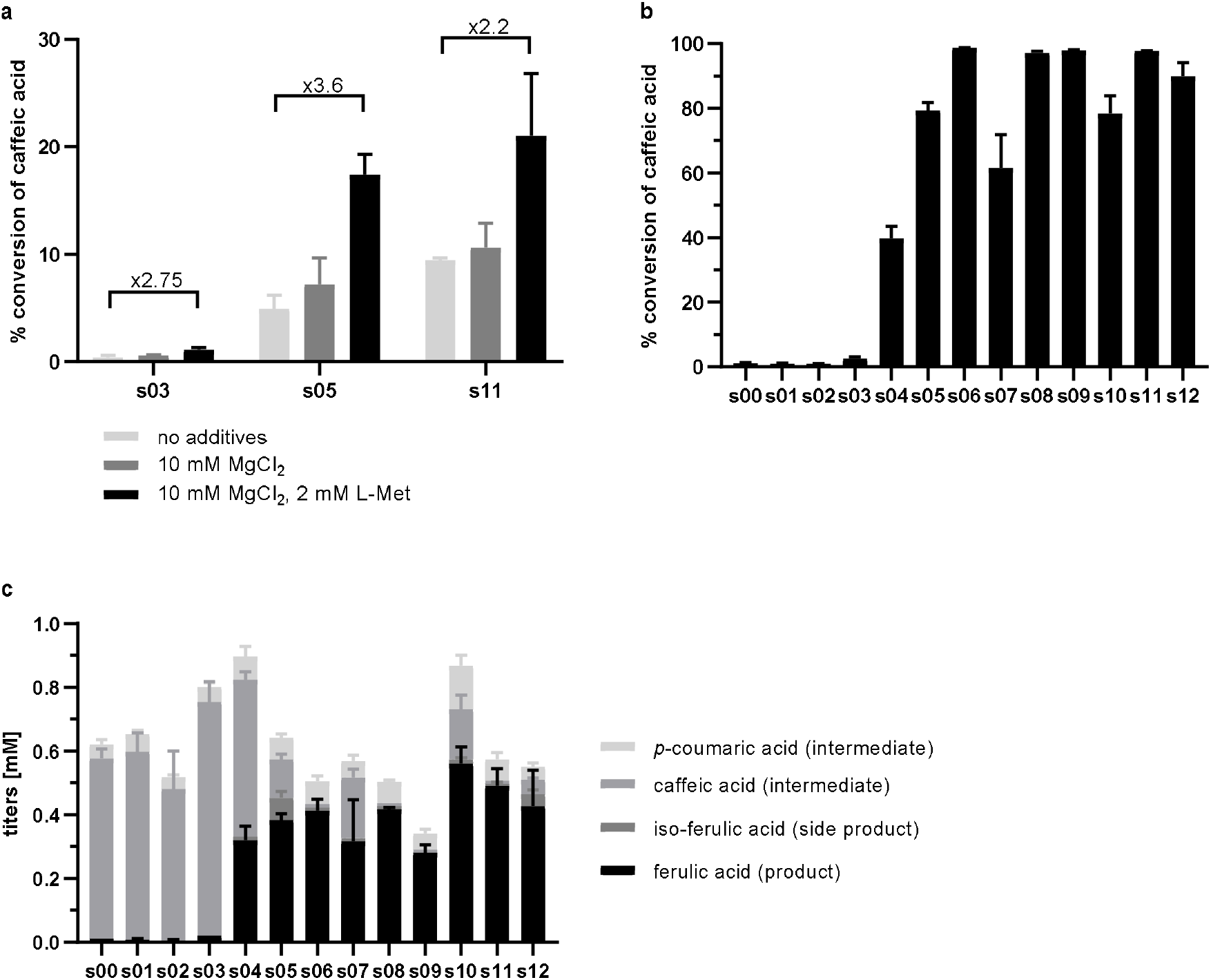
Fermentation of *E. coli* K12 MG1655(DE3) expressing a recombinant pathway to produce(iso-)ferulic acid. a) Media optimization performed with feeding of the pathway intermediate caffeic acid (2 mM); data displayed as percent conversion of fed caffeic acid. b) and c) Experiment performed with optimized media with glucose as a carbon source and L-Tyrosine (3mM) as a pathway precursor; data displayed as percent conversion of produced caffeic acid (panel b) or as stacked histogram of titers of product, side product and pathway intermediates (panel c). Mean +/− SD, n=3.

## Discussion

Closing the gap between computational annotations and the biotechnological exploitation of natural enzymes as industrial biocatalysts requires extensive functional screening of enzyme libraries. Alternatively, scarce sampling of enzyme families and deep functional analysis (“thick data”) can be utilized to improve annotation pipelines and thus the interpretation of big data. In this study, we developed a rapid prototyping platform for SAM-dependent methyltransferases, an enzyme superfamily that has great potential for functionalization of natural product-inspired pharmaceuticals. The chosen approach follows in the footsteps of a range of studies utilizing *in vitro* transcription/translation systems for prototyping of antibodies^43^, protein expression enhancing factors^44^, transcription regulatory elements^45^, GPCRs^46^, quorum-sensing systems^47^ and entire biosynthetic pathways^48^, which highlights the generalizability of this expression system^49^. In our study we observed that the enzymatic activity is highly reproducible across biological replicates of the *in vitro* transcription/translation system, which is in good agreement with the previous studies. In addition to the time and effort saved on molecular cloning, transformation and protein expression in *E. coli* (1-3 weeks), another advantage is that no additional lysis and clearing steps are required before the enzymatic reaction is performed. This is particularly advantageous for enzymes requiring cofactors that cannot cross the cell membrane, such as SAM. In our workflow, we combined the *in vitro* expression system with a microliter-scale enzymatic assay coupled to a TR-FRET read-out. While this read-out is sensitive to experimental error due to the small volumes and the required dilution steps, we were able to generate robust results by using master mixes and multichannel pipettes for all steps. The TR-FRET-based detection of the SAM cofactor - rather than a specific substrate or product - allows for the screening of a library of substrates. The format can furthermore be used to swiftly optimize reaction conditions, such as buffers, salts and substrate concentrations and thus generate “thick data”. The entire workflow should be amenable to automation by using liquid handling robots and is therefore scalable to also screen large enzyme libraries.

The workflow allowed us to rapidly screen ~30 enzymes against 5 substrates. In particular for caffeic acid as a substrate, we identified several distant homologues with remarkable activity, two of which don’t even carry the conserved active site residues. With a more conservative *in silico* approach of selecting enzymes of interest, we might not even have considered these as suitable enzymes. However, our approach allowed us to cast a wide net, explore the activity of these distantly related enzymes and use them in an *E. coli* cell factory for ferulic acid, a methylated phenolic acid of pharmaceutical interest. Zooming in on these unexpected hits and their close relatives with structural and functional studies will allow us to better understand the underlying mechanisms of substrate selectivity and regioselectivity in OMTs.

Lastly, we tested a subset of the pre-screened OMTs in the context of a recombinant biosynthetic pathway in *E. coli*. We observed that all enzymes seen to be active in the pre-screening step were also catalytically active in the pathway, whereas enzymes found to be inactive in the screen, remained inactive in the pathway. The trends in substrate conversion levels and regio-selectivity, however, were not necessarily correlated between the *in vitro* and in vivo experiments. This is a hurdle well known to metabolic engineers and is inherent to *in vitro* characterization of enzymes. However, the high cost of *in vivo* screening in terms of time and consumables justifies the need for *in vitro* prototyping.

## Methods

### Selection of enzymes of interest

Two different multiple sequence alignments of known OMTs were generated with the Clustal Omega EBI webtool^22^ (Extended Data Table 1) and used as input for HMMsearch (EBI webtool version 2.23.0^21^; data base of reference proteomes of all taxa excluding green plants (taxid: 33090), significance E-value cut-off 0.01 for the entire sequence and 0.03 for hits). The significant results were combined into one data set (Supplementary File 1) and used as an input for calculating a sequence similarity network with a webtool of the Enzyme Function Initiative (EFI-EST^29^; node selection cut-off: protein length between 180 and 400 amino acids, edge selection cut-off: alignment score >30). The finalized network was visualized in Cytoscape 3.8.0^30^ with the yFiles organic layout. For the representation in Extended Data Figure 1, the nodes were further filtered to exclude all nodes with an HMM score below 70 and all edges with sequence identity below 50%. From the thus generated clusters enzymes were chosen for experimental characterization (Table 1).

### Design and synthesis of DNA templates for TXTL reactions

The selected genes were codon-optimized for expression in *E. coli* with the Integrated DNA Technologies (IDT) optimization algorithm and manually modified to exclude recognition sites for BsaI, NcoI, XhoI and where possible NdeI restriction enzymes. The 5’ end of all DNA fragments was designed with an overhang of 500bp, the p70a promoter sequence and an NcoI recognition site to facilitate cloning into the pET21b(+) (Novagen) and pBEST (Arbor Bioscience) expression vectors. The 3’ end was designed to include an XhoI recognition site, the T500 terminator and a 500bp overhang. The synthetic DNA was obtained from Arbor Bioscience (Ann Arbor, MI, USA) with an additional purification step to allow for direct use in the myTXTL^®^ reaction for linear templates.

### Construction of plasmids

All molecular cloning and plasmid propagation steps were performed in chemically competent *Escherichia coli E. cloni*^®^ 10G (F-*mcrA* Δ(*mrr-hsd*RMS-*mcr*BC) *end*A1 *rec*A1 Φ80*dlac*ZΔM15 Δ*lac*X74 *ara*D139 Δ(*ara,leu*)7697*gal*U *gal*K *rps*L *nup*G λ-*ton*A) produced by Lucigen (Middleton, WI, USA). Genes encoding for OMTs selected for *in vivo* testing were cloned directly from the synthetic DNA fragments by restriction and ligation (NcoI/XhoI) into pET21b(+) for expression under the T7 promoter (Extended Data Table 2). From there the genes were amplified by polymerase chain reaction (PCR) with gene specific 5’primers and the T7 terminator primer to generate an NdeI recognition site at the 5’ prime end. The PCR products were inserted by restriction and ligation (NdeI/XhoI) into the second multiple cloning site of the plasmid c71 (pRSF::FjTAL) for expression under the T7 promoter (Extended Data Table 2). All constructs were verified by sequencing by ETON Bioscience (Charlestown, MA, USA). Plasmids c71, c84 and c86 were constructed in a previous study^31^.

### **In vitro** transcription/translation

*In vitro* transcription/translation was performed with the myTXTL^®^ kit from Arbor Bioscience according to the manufacturer’s instructions. In brief, the synthesized DNA fragments were dissolved in nuclease-free water to a final concentration of 109.1 nM and stored at −70°C between experiments. All assay components were thawed on ice (myTXTL^®^ lysate, GamS protein and DNA templates) and mixed by carefully pipetting up and down. To minimize pipetting errors, a master mix of myTXTL^®^ lysate (9μL per reaction) and GamS protein (0.8 μL per reaction) was prepared on ice and aliquoted into 1.5 mL microcentrifuge tubes. 2.2 μL of DNA template were added to each tube and mixed by carefully pipetting up and down (final concentration 20 nM). The reactions were incubated on ice for 5 min and then transferred to a water bath at 29°C for 16 h. As negative controls (“no OMT”), one reaction was performed with a DNA template not encoding for an OMT enzyme and one reaction with nuclease-free water without DNA.

### SDS PAGE

To visualize protein expression 1 μL of the TXTL reactions was mixed with 2 μL of water and 3 μL of 2x Laemmli loading dye (Bio-Rad, Hercules, CA, USA) and incubated at 90°C for 3 min. The denatured samples were loaded onto AnyKD™ Mini-PROTEAN^®^ TGX™ precast protein gels (Bio-Rad, Hercules, CA, USA) and separated for 40 min at 40 mA. Protein bands were visualized by staining with InstantBlue^®^ protein stain and imaging with the Bio-Rad ChemiDoc™ imager.

### Expression and purification of MxSafC

Plasmid c157 was transformed into chemically competent *E. coli* BL21 DE3 (F^−^ *ompT gal dcm lon hsdS*_*B*_(*r*_*B*_^−^ *m*_*B*_^−^) λ(DE3 [*lacI lacUV5-T7p07 ind1 sam7 nin5*]) [*malB*+]_K-12_(λ^S^)) and maintained on selective LB agar containing 100 μg/mL carbenicillin. A starter culture was inoculated from a single colony (5 mL, LB with carbenicillin) and incubated overnight at 37°C, 250 rpm. The main culture was inoculated from the starter culture (1:100) and incubated at 37°C, 250 rpm until an optical density OD_600_ of 0.7 was reached. Expression was induced with Isopropyl β-D-1-thiogalactopyranoside (IPTG, 1 mM final) and the temperature was lowered to 30°C (250 rpm, overnight). All following steps were performed at 4°C with chilled buffers. The cells were harvested by centrifugation (10 min, 3,000 rpm) and resuspended in 20 mL lysis buffer (buffer A including one EDTA-free protease inhibitor tablet (Roche) and 10 mg/mL lysozyme; buffer A: 50 mM Tris/HCl pH 7.4, 500 mM NaCl, 10 mM imidazole). The cell suspension was incubated on ice for 20 min, lysed by sonication (20% duty cycle, 10 cycles of 15 s ON/15 s OFF) and cleared by centrifugation for 20 min at 40,000 x g. The supernatant was loaded onto the affinity matrix equilibrated with buffer A by gravity flow (Qiagen, Ni-NTA agarose slurry, 0.25 mL column volume). The column was washed with 20 column volumes of buffer A and eluted stepwise with one column volume of buffers B1 to B6 (buffers B1-B6: 50 mM Tris/HCl pH 7.4, 500 mM NaCl, 50 mM imidazole/ 100 mM imidazole/ 150 mM imidazole/ 200 mM imidazole/ 250 mM imidazole or 500 mM imidazole, respectively). The eluates of each step were collected in separate fractions and analyzed by SDS PAGE. MxSafC containing fractions with low protein background were pooled and dialyzed overnight at 4°C against storage buffer (20 mM Tris/HCl pH 7.4, 50 mM NaCl, 0.2 mM MgCl_2_, 2 mM DTT). The protein concentration was determined by absorbance at 280 nm (NanoDrop, ThermoFisher Scientific, USA) before the purified enzyme was aliquoted and stored at −70°C.

### **In vitro** OMT reaction

The *In vitro* OMT reaction was adapted from the conditions used by Siegrist et al.^32^. To minimize pipetting errors, a master mix including all reaction components, but the enzyme was prepared (50 mM HEPES/NaOH pH 7, 20 mM MgCl_2_, 2 mM SAM, 2 mM substrate (from 40x stock in DMSO)). The total reaction volume was 42 μL, with 2 μL of purified enzyme/ TXTL reaction used in the initial experiment and later 5 μL of a diluted stock (2.5-fold dilution in OMT reaction buffer) to further minimize pipetting errors. After aliquoting the master mix into 96-well microtiter plates (200 μL round-bottom plates), the TXTL samples were added and mixed by carefully pipetting up and down. Purified MxSafC enzyme was included in one well as a positive control. The “no OMT” controls (see section “*in vitro* transcription/translation”) were treated like the other enzyme samples. The sealed plates were incubated at 30°C for 24 h before the reactions were quenched with HClO_4_ (final 2% v/v from a 10% v/v stock) and centrifuged. The supernatants were analyzed by Time Resolved-Fluorescence Energy Resonance Transfer (TR-FRET) and (optionally) by High Performance Liquid Chromatography (HPLC).

### TR-FRET assay for SAM detection

To detect the consumption of the SAM co-factor as a measure of OMT reactivity, we used the TR-FRET Bridge-It^®^ S-Adenosyl Methionine (SAM) Fluorescence Assay Kit from Mediomics LLC (St. Louis, Missouri) according to the manufacturer’s instructions with slight modifications. In brief, we thawed the assay solution at 37°C for 30 min and transferred 18 μL into the wells of a white 384-well round-bottom polystyrene plate (Corning, NY, USA). We diluted the quenched OMT reactions 21-fold by mixing 2 μL of the reaction with 40 μL of water by pipetting up and down, and transferred the samples to the 384-well plate without bubbling. In addition to the “no OMT” controls (no SAM consumption expected), one or two wells were measured with only the TR-FRET assay solution (20 μL, “blank”). The plate was incubated in the dark for 30 min at room temperature before measuring the TR-FRET signal in a Tecan Infinite-200 plate reader with the following settings: *mode*: fluorescence top reading, *excitation wavelength*: 340 nm, *emission wavelength*: 667 nm, *excitation bandwidth*: 9 nm, *emission bandwidth*: 20 nm, *gain*: 220 (manual), *number of flashes*: 100, *integration time*: 400 μs, *lag time*: 50 μs, *settle time*: 150 ms. The ratio of the acceptor channel counts to the donor channel counts was calculated for all measured wells (FRET), baseline corrected with the FRET ratio of the “blank” and normalized to the average of the FRET ratio of the “no OMT” controls to obtain the relative SAM detection (Eq. 1). In the initial experiment, the samples were handled with single-channel pipettes, whereas in the later experiments, multichannel pipettes were used throughout to minimize pipetting errors.

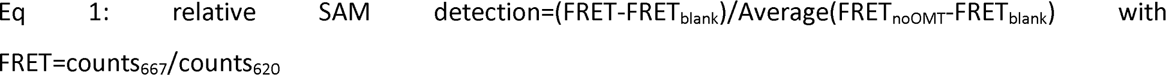

### Fermentation

The OMT encoding plasmids were transformed into chemically competent^33^ *E. coli* K12 MG1655(DE3)^34^ already bearing the plasmids c84 and c86 encoding for CYP199A2 F185L NΔ7 and its redox partners putidaredoxin (Pux) and putidaredoxin reductase (PuR). All strains generated in this way are listed in Extended Data Table 3. The final strains were maintained on selective media with carbenicillin, spectinomycin and kanamycin at all times. Starter cultures were prepared from three individual colonies of the final strains in 5 mL Lysogeny broth (LB) supplemented with carbenicillin (100 μg/mL), spectinomycin (50 μg/mL) and kanamycin (50 μg/mL) in round-bottom polystyrene tubes, incubated over night at 37°C with agitation and used to inoculate the main cultures (7 mL LB with antibiotics; round-bottom polystyrene tubes). After 4 h of growth at 37°C, 250 rpm, OD_600_ was measured and the appropriate volume of each culture pelleted and resuspended in modified, selective M9 medium including substrates and 4% glucose to obtain 15 mL cultures at OD_600_ of 0.7 in sterile glass tubes. These cultures were incubated at 26°C, 160 rpm for 96 h. Samples of 200 μL were taken after 96 h and quenched with 50 μL of HClO_4_ (10 % (v/v) stock), spun for 10 min at 20,000 x g and the supernatants were analyzed by HPLC. Media optimization was performed in small scale (5 mL in round-bottom polystyrene tubes) throughout the entire experiment.

M9 medium composition (1x) prepared from sterile stocks: M9 salts (Millipore-Sigma, used as 5x stock), Trace Mineral Supplement (ATCC^®^ MD-TMS™, used as 200x stock), vitamin mix (from 100x stock; final: riboflavin 0.84 mg/L, folic acid 0.084 mg/L, nicotinic acid 12.2 mg/L, pyridoxine 2.8 mg/L, and pantothenic acid 10.8 mg/L), biotin (from 1000x stock; final: 0.24 mg/L), thiamine (from 1470x stock; final: 340 mg/L), δ-Aminolevulinic acid (from 1000x stock in MeOH, final: 7.5 μg/mL), IPTG (from 1000x stock, final: 1 mM), carbenicillin (from 1000x stock, final: 100 μg/mL), spectinomycin (from 1000x stock, final: 50 μg/mL), kanamycin (from 1000x stock, final: 50 μg/mL, 4% (w/v) glucose (from 50% w/v stock). Additives for media optimization experiments: caffeic acid (from fresh 100x stock in MeOH, final 2 mM) and either a) no further additives, b) MgCl_2_ (from 500x sterile stock in water, final 2 mM) or c) MgCl_2_ (from 500x sterile stock in water, final 2 mM) and L-methionine (from fresh 100x stock in 1M HCl, final 10 mM). Additives for all other experiments: MgCl_2_ (from 500x sterile stock in water, final 2 mM) and L-methionine and L-tyrosine (from fresh joined 100x stock in 1M HCl, final 10 mM and 3 mM, respectively).

### HLPC analysis

The supernatants of the quenched *in vitro* OMT reactions and fermentation samples were analyzed by reversed-phase HPLC (instrument: Agilent 1100; autosampler: HiP sampler G1367A, T=4°C, 10 μL injection; column: Agilent Zorbax Eclipse XDB-C18 80Å, 4.6 x 150 mm, 5μm, T=30°C; detector: Agilent diode array detector G1315B, λ=275 nm (catechol and methylated products) and λ=310nm ((iso-)ferulic acid and pathway intermediates); gradient: 10% to 35% Acetonitrile with 0.1% Trifluoracetic acid over 17 min). The peaks for products and intermediates were identified by comparing the retention times to authentic standards. The integrated peak areas were converted to concentrations in mM based on calibration curves generated with authentic standards.

### Sequence analysis of putative OMTs

We aligned the sequences with mafft v7.310^35^b (--genafpair), inferred the maximum likelihood phylogenies with FastTree v2.1.10^36^ and visualized the tree and the corresponding activity heat map in Figure 4 with the R packages ggplot2^37^ and ggtree^38^.

## Acknowledgements

KH is grateful for the support by the Human Frontier Science Program (Grant Number LT000969/2016-L). This work was supported by the MIT Portugal Program (Grant Number 6937822).

## Author Contributions

KH conceived the study, selected the putative OMTs, performed all experiments and wrote the manuscript with support and guidance by K.L.J.P. KH and TH analyzed the data and created figures. TH provided bioinformatics support. All authors read and approved the final version of this manuscript.

## Extended Data Tables

**Extended Data Table 1:**
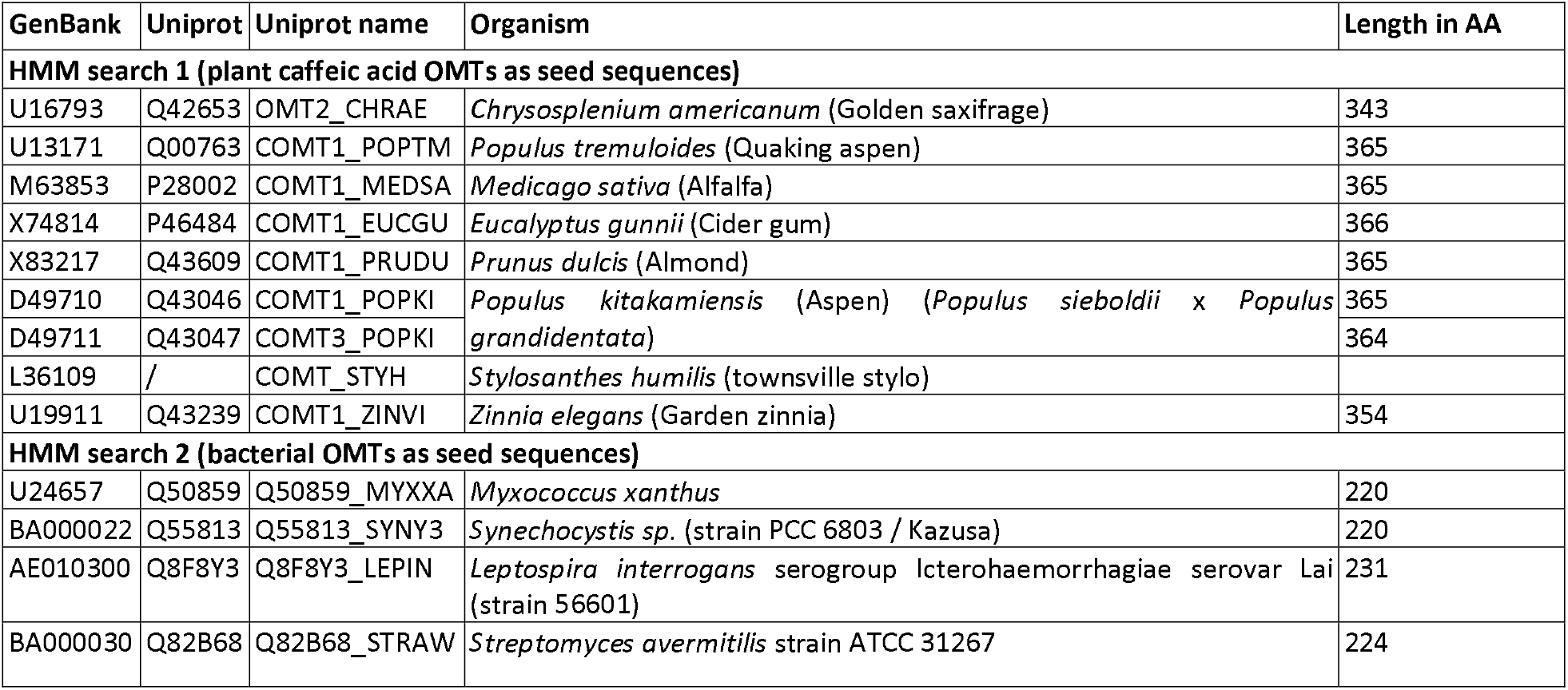
Table of input sequences for HMMsearch1 and 2.

**Extended Data Table 2:**
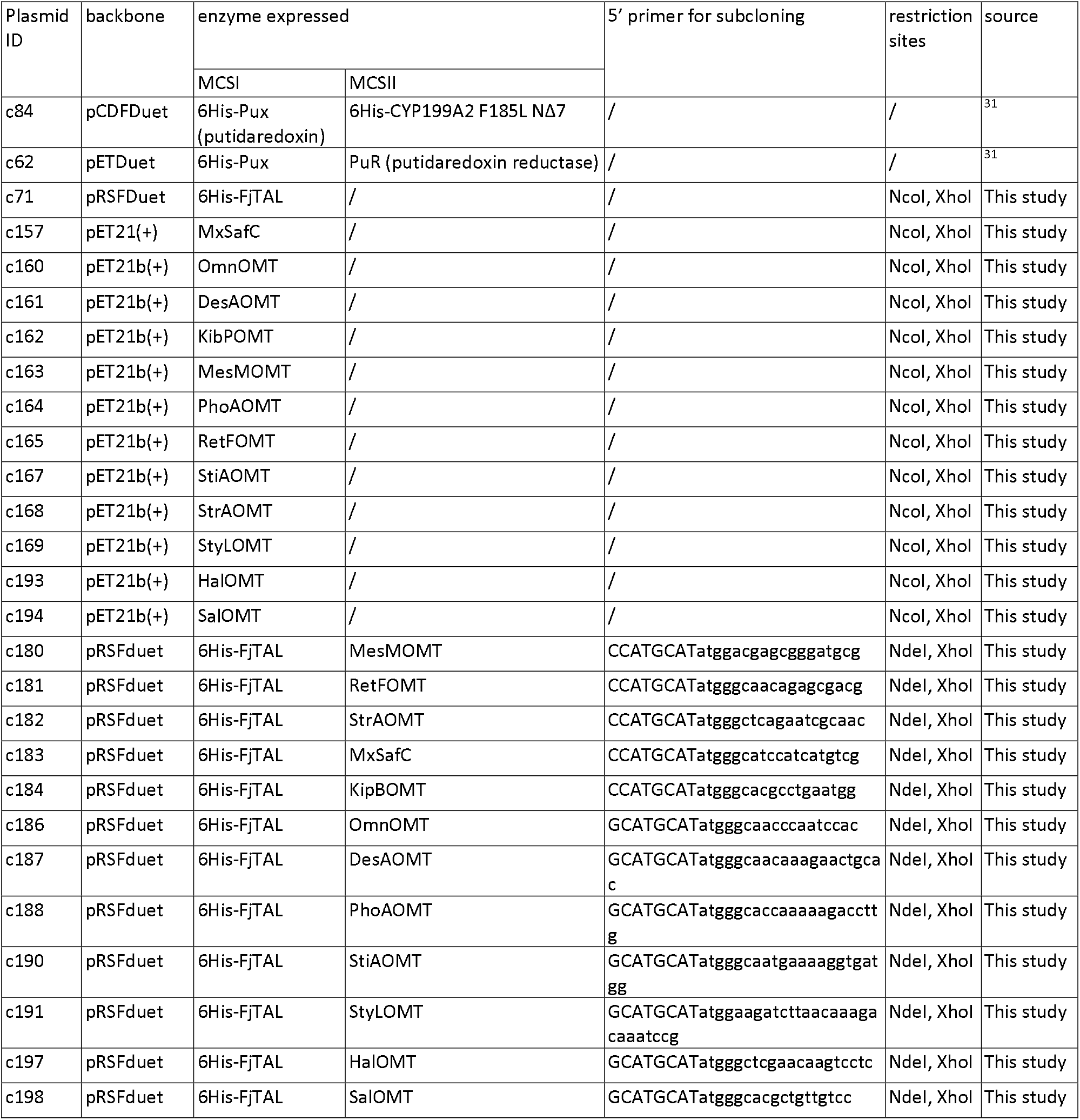
List of plasmids used in this study with the enzymes expressed from the respective multiple cloning sites (MCS), the 5’ primer sequences and the restriction sites used for subcloning.

**Extended Data Table 3:** List of bacterial strains used in this study to produce (iso-)ferulic acid from fed L-tyrosine or caffeic acid.

| Strain ID | plamids | enzymes expressed | source |
| --- | --- | --- | --- |
| s00 | c84<br>c62<br>c71 | 6His-Pux, 6His-CYP199A2F185L NΔ7<br>6His-Pux, PuR<br>FjTAL | <sup>31</sup> |
| s01 | c84<br>c62<br>c197 | 6His-Pux, 6His-CYP199A2F185L NΔ7<br>6His-Pux, PuR<br>FjTAL, HalOMT | This study |
| s02 | c84<br>c62<br>c198 | 6His-Pux, 6His-CYP199A2F185L NΔ7<br>6His-Pux, PuR<br>FjTAL, SalOMT | This study |
| s03 | c84<br>c62<br>c180 | 6His-Pux, 6His-CYP199A2F185L NΔ7<br>6His-Pux, PuR<br>FjTAL, MesMOMT | This study |
| s04 | c84<br>c62<br>c187 | 6His-Pux, 6His-CYP199A2F185L NΔ7<br>6His-Pux, PuR<br>FjTAL, DesAOMT | This study |
| s05 | c84<br>c62<br>c183 | 6His-Pux, 6His-CYP199A2F185L NΔ7<br>6His-Pux, PuR<br>FjTAL, MxSafC | This study |
| s06 | c84<br>c62<br>c188 | 6His-Pux, 6His-CYP199A2F185L NΔ7<br>6His-Pux, PuR<br>FjTAL, PhoAOMT | This study |
| s07 | c84<br>c62<br>c184 | 6His-Pux, 6His-CYP199A2F185L NΔ7<br>6His-Pux, PuR<br>FjTAL, KipBOMT | This study |
| s08 | c84<br>c62<br>c191 | 6His-Pux, 6His-CYP199A2F185L NΔ7<br>6His-Pux, PuR<br>FjTAL, StyLOMT | This study |
| s09 | c84<br>c62<br>c181 | 6His-Pux, 6His-CYP199A2F185L NΔ7<br>6His-Pux, PuR<br>FjTAL, RetFOMT | This study |
| s10 | c84<br>c62<br>c190 | 6His-Pux, 6His-CYP199A2F185L NΔ7<br>6His-Pux, PuR<br>FjTAL, StiAOMT | This study |
| s11 | c84<br>c62<br>c182 | 6His-Pux, 6His-CYP199A2F185L NΔ7<br>6His-Pux, PuR<br>FjTAL, StrAOMT | This study |
| s12 | c84<br>c62<br>c186 | 6His-Pux, 6His-CYP199A2F185L NΔ7<br>6His-Pux, PuR<br>FjTAL, OmnOMT | This study |

**Extended Data Table 4:** Regioselectivity of OMT enzymes in vitro (pre-screen) and in vivo (fermentation) expressed as regio-isomeric excess of the meta-over the para-product (RE=(c[meta]-c[para])/(c[meta]+c[para])*100).

| OMT enzyme | pre-screen |  | fermentation |  |
| --- | --- | --- | --- | --- |
|  | RE mean | SD | RE mean | SD |
| HalOMT | n.d. | n.d. | n.d. | n.d. |
| SalOMT | n.d. | n.d. | n.d. | n.d. |
| MesMOMT | 86.44 | 0.00 | 100.00 | 0.00 |
| DesAOMT | 32.09 | 7.01 | 93.87 | 0.76 |
| MxOMT | -34.78 | 1.17 | 69.24 | 7.80 |
| PhoAOMT | 39.04 | 6.29 | 95.25 | 0.39 |
| KibpOMT | 46.40 | 5.92 | 93.13 | 2.88 |
| StyLOMT | 49.06 | 3.81 | 95.31 | 0.06 |
| RetFOMT | 88.04 | 9.95 | 100.00 | 0.00 |
| StiAOMT | 48.83 | 9.18 | 95.41 | 1.64 |
| StrAOMT | 62.38 | 9.45 | 97.27 | 2.41 |
| OmnOMT | -2.96 | 3.41 | 84.65 | 2.58 |

## Extended Data Figures

**Extended Data Figure 1:**
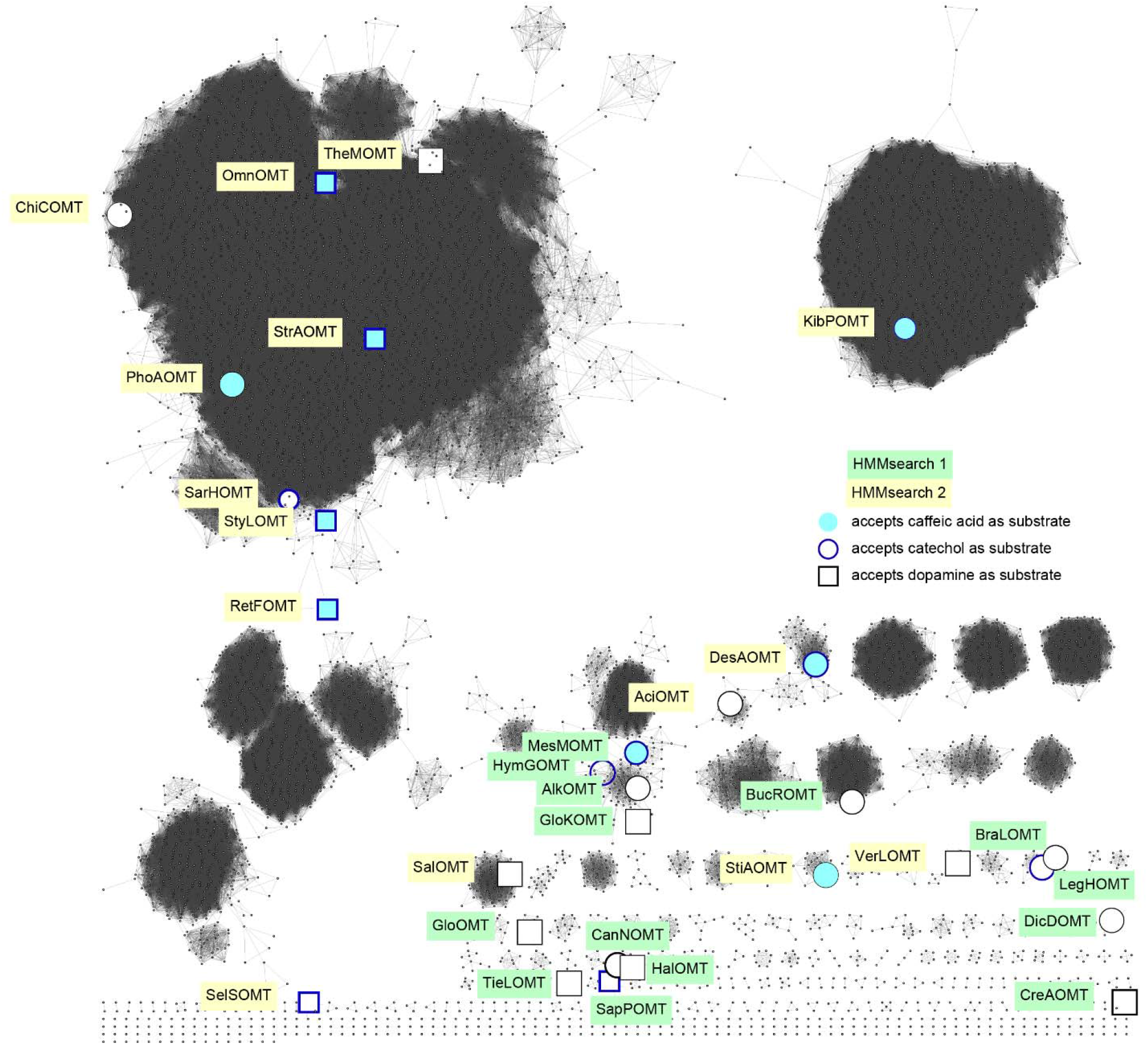
Sequence similarity network calculated with the EFI-EST webtool and visualized with the yFiles organic layout in Cytoscape with an edge cut-off at 50% sequence identity. Only nodes with an HMMsearch score of 70 or higher were included in the generation of the network. Enlarged nodes with label: enzymes tested in this study; Yellow box: enzymes with high sequence similarity to the bacterial input sequences, green box: enzymes with high sequence similarity to the plant input sequences; open black circle: inactive on tested substrates, filled symbol: active on caffeic acid, blue outline: active on catechol, rectangle: active on dopamine.

**Extended Data Figure 2:**
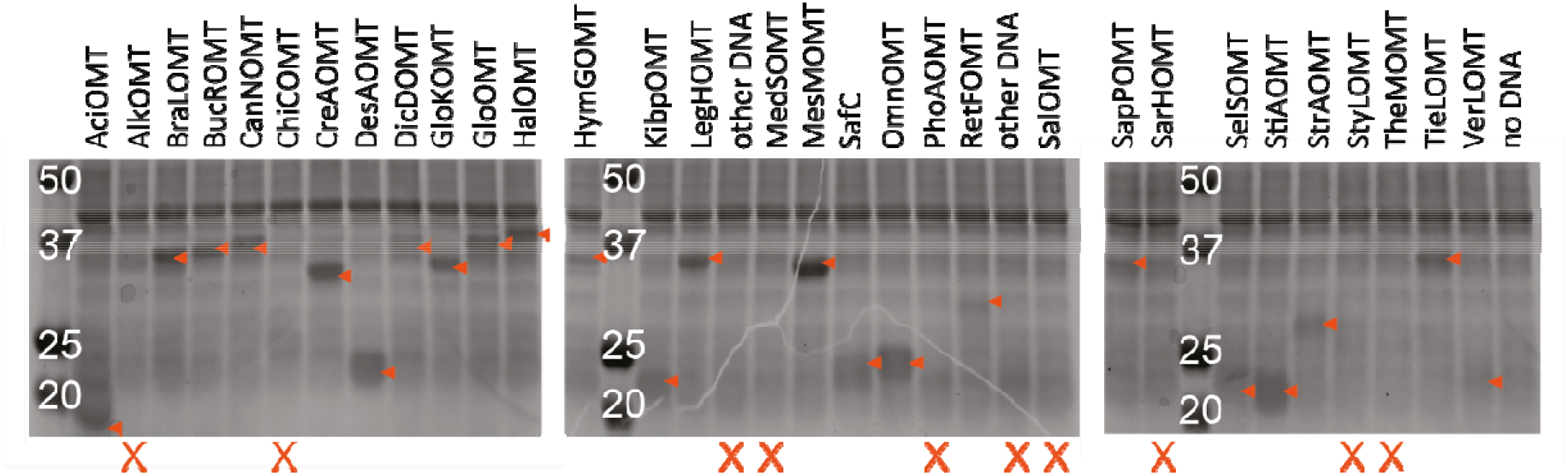
Coomassie stained SDS PAGE of putative OMTs expressed in TXTL reactions. Orange arrows indicate protein bands suspected to represent the respective OMT. Lanes without obvious bands of the correct molecular weight are marked with X below.

## References

1. Liscombe, D. K., Louie, G. V. & Noel, J. P. Architectures, mechanisms and molecular evolution of natural product methyltransferases. Natural Product Reports 29, 1238 (2012).

2. Vanholme, R., Demedts, B., Morreel, K., Ralph, J. & Boerjan, W. Lignin biosynthesis and structure. Plant physiology 153, 895–905 (2010).

3. Kadoma, Y. & Fujisawa, S. A comparative study of the radical-scavenging activity of the phenolcarboxylic acids caffeic acid, p-coumaric acid, chlorogenic acid and ferulic acid, with or without 2-mercaptoethanol, a thiol, using the induction period method. Molecules 13, 2488–2499 (2008).

4. Taram, F., Winter, A. N. & Linseman, D. A. Neuroprotection comparison of chlorogenic acid and its metabolites against mechanistically distinct cell death-inducing agents in cultured cerebellar granule neurons. Brain Research 1648, 69–80 (2016).

5. Koirala, N., Thuan, N. H., Ghimire, G. P., Thang, D. Van & Sohng, J. K. Methylation of flavonoids: Chemical structures, bioactivities, progress and perspectives for biotechnological production. Enzyme and Microbial Technology 86, 103–116 (2016).

6. Wen, L. et al. Structure, bioactivity, and synthesis of methylated flavonoids. Annals of the New York Academy of Sciences 1398, 120–129 (2017).

7. Li, Y. et al. Dissociation of Antimicrobial and Hemolytic Activities of Gramicidin S through N-Methylation Modification. ChemMedChem 8, 1865–1872 (2013).

8. Das, D. et al. Synthesis, SAR and biological studies of sugar amino acid-based almiramide analogues: N-methylation leads the way. Org. Biomol. Chem 15, 3337 (2017).

9. Zahid Khan, M. & Nawaz, W. The emerging roles of human trace amines and human trace amine-associated receptors (hTAARs) in central nervous system. Biomedicine and Pharmacotherapy 83, 439–449 (2016).

10. Finkle, B. J. & Nelson, R. F. Enzyme reactions with phenolic compounds: a meta-O-methyltransferase in plants. Biochimica et Biophysica Acta 78, 747–749 (1963).

11. Higuchi, T., Shimada, M., Nakatsubo, F. & Tanahashi, M. Differences in Biosyntheses of Guaiacyl and Syringyl Lignins in Woods. Wood Science and Technology 11, (Springer-Verlag, 1977).

12. Hicks, M. a. & Prather, K. L. J. Bioprospecting in the Genomic Age. Advances in Applied Microbiology 87, (Elsevier Inc., 2014).

13. Chao, R., Mishra, S., Si, T. & Zhao, H. Engineering biological systems using automated biofoundries. doi:10.1016/j.ymben.2017.06.003

14. Casini, A. et al. A Pressure Test to Make 10 Molecules in 90 Days: External Evaluation of Methods to Engineer Biology. J Am Chem Soc 140, 4302–4316 (2018).

15. Jacques, P. et al. High-throughput strategies for the discovery and engineering of enzymes for biocatalysis. Bioprocess and Biosystems Engineering 40, 161–180 (2017).

16. Longwell, C. K., Labanieh, L. & Cochran, J. R. High-throughput screening technologies for enzyme engineering. doi:10.1016/j.copbio.2017.05.012

17. Garamella, J., Marshall, R., Rustad, M. & Noireaux, V. The All E. coli TX-TL Toolbox 2.0: A Platform for Cell-Free Synthetic Biology. ACS Synthetic Biology 5, 344–355 (2016).

18. Shin, J. & Noireaux, V. Efficient cell-free expression with the endogenous E. coli RNA polymerase and sigma factor 70. Journal of Biological Engineering 4, 8 (2010).

19. Heyduk, T. & Heyduk, E. Molecular beacons for detecting DNA binding proteins. Nature Biotechnology 20, 171–176 (2002).

20. Tian, L., Wang, R. E., Fei, Y. & Chang, Y.-H. A homogeneous fluorescent assay for cAMP-phosphodiesterase enzyme activity. Journal of biomolecular screening 17, 409–14 (2012).

21. Potter, S. C. et al. HMMER web server: 2018 update. Nucleic Acids Research 46, W200–W204 (2018).

22. Sievers, F. et al. Fast, scalable generation of high-quality protein multiple sequence alignments using Clustal Omega. Molecular systems biology 7, 539 (2011).

23. Joshi, C. P. & Chiang, V. L. Conserved sequence motifs in plant S-adenosyl-L-methionine-dependent methyltransferases. Plant Molecular Biology 37, 663–674 (1998).

24. Liu, X. et al. Systematic analysis of O-methyltransferase gene family and identification of potential members involved in the formation of O-methylated flavonoids in Citrus. Gene 575, 458–472 (2016).

25. Nelson, J. T., Lee, J., Sims, J. W. & Schmidt, E. W. Characterization of SafC, a catechol 4-O-methyltransferase involved in saframycin biosynthesis. Applied and Environmental Microbiology 73, 3575–3580 (2007).

26. Kopycki, J. G. et al. Functional and structural characterization of a cation-dependent O-methyltransferase from the cyanobacterium [i]Synechocystis[/i] sp. strain PCC 6803. Journal of Biological Chemistry 283, 20888–20896 (2008).

27. Hou, X. et al. Crystal structure of SAM-dependent O-methyltransferase from pathogenic bacterium Leptospira interrogans. Journal of Structural Biology 159, 523–528 (2007).

28. Youngdae, Y. et al. Characterization of an O-methyltransferase from [i]Streptomyces avermitilis[/i] MA-4680. Journal of Microbiology and Biotechnology 20, 1359–1366 (2010).

29. Gerlt, J. A. et al. Enzyme Function Initiative-Enzyme Similarity Tool (EFI-EST): A web tool for generating protein sequence similarity networks. Biochimica et biophysica acta 1854, 1019–37 (2015).

30. Shannon, P. et al. Cytoscape: A Software Environment for Integrated Models. Genome Research 13, 2498–504 (2003).

31. Haslinger, K. & Prather, K. L. J. Heterologous caffeic acid biosynthesis in Escherichia coli is affected by choice of tyrosine ammonia lyase and redox partners for bacterial Cytochrome P450. Microbial Cell Factories 19, (2020).

32. Siegrist, J. et al. Functional and structural characterisation of a bacterial O-methyltransferase and factors determining regioselectivity. FEBS Letters 591, 312–321 (2017).

33. Inoue, H., Nojima, H. & Okayama, H. High efficiency transformation of Escherichia coli with plasmids. Gene 96, 23–28 (1990).

34. Nielsen, D. R., Yoon, S.-H., Yuan, C. J. & Prather, K. L. J. Metabolic engineering of acetoin and meso-2, 3-butanediol biosynthesis in E. coli. Biotechnology journal 5, 274–84 (2010).

35. Nakamura, T., Yamada, K. D., Tomii, K. & Katoh, K. Parallelization of MAFFT for large-scale multiple sequence alignments. Bioinformatics 34, 2490–2492 (2018).

36. Price, M. N., Dehal, P. S. & Arkin, A. P. FastTree 2 - Approximately maximum-likelihood trees for large alignments. PLoS ONE 5, e9490 (2010).

37. Wickham, H. ggplot2. Wiley Interdisciplinary Reviews: Computational Statistics 3, 180–185 (2011).

38. Yu, G., Tsan, T., Lam, Y., Zhu, H. & Guan, Y. Two Methods for Mapping and Visualizing Associated Data on Phylogeny Using Ggtree. Mol. Biol. Evol. 3 35, 3041–3043 (2018).

39. Zubieta, C., Kota, P., Ferrer, J., Dixon, R. a & Noel, J. P. Structural Basis for the Modulation of Lignin Monomer Methylation by Caffeic Acid / 5-Hydroxyferulic Acid 3 / 5-O-Methyltransferase. The Plant Cell 14, 1265–1277 (2002).

40. Brandt, W., Manke, K. & Vogt, T. A catalytic triad - Lys-Asn-Asp - Is essential for the catalysis of the methyl transfer in plant cation-dependent O-methyltransferases. Phytochemistry 113, 130–139 (2015).

41. Rodrigues, J. L., Araújo, R. G., Prather, K. L. J., Kluskens, L. D. & Rodrigues, L. R. Heterologous production of caffeic acid from tyrosine in Escherichia coli. Enzyme and Microbial Technology 71, 36–44 (2015).

42. Kunjapur, A. M., Hyun, J. C. & Prather, K. L. J. Deregulation of S-adenosylmethionine biosynthesis and regeneration improves methylation in the E. coli de novo vanillin biosynthesis pathway. Microbial Cell Factories 15, 61 (2016).

43. Yin, G. et al. Aglycosylated antibodies and antibody fragments produced in a scalable in vitro transcription-translation system. mAbs 4, 217–225 (2012).

44. Woodrow, K. A. & Swartz, J. R. A sequential expression system for high-throughput functional genomic analysis. PROTEOMICS 7, 3870–3879 (2007).

45. McManus, J. B., Emanuel, P. A., Murray, R. M. & Lux, M. W. A method for cost-effective and rapid characterization of engineered T7-based transcription factors by cell-free protein synthesis reveals insights into the regulation of T7 RNA polymerase-driven expression. Archives of Biochemistry and Biophysics 674, 108045 (2019).

46. Cortès, S., Hibti, F. E., Chiraz, F. & Ezzine, S. High-throughput E. coli cell-free expression: From PCR product design to functional validation of GPCR. in Methods in Molecular Biology 2025, 261–279 (Humana Press Inc., 2019).

47. Halleran, A. D. & Murray, R. M. Cell-Free and In Vivo Characterization of Lux, Las, and Rpa Quorum Activation Systems in E. coli. ACS Synthetic Biology 7, 752–755 (2018).

48. Dudley, Q. M., Nash, C. J. & Jewett, M. C. Cell-free biosynthesis of limonene using enzyme-enriched Escherichia coli lysates. Synthetic Biology 4, (2019).

49. Dopp, J. L., Rothstein, S. M., Mansell, T. J. & Reuel, N. F. Rapid prototyping of proteins: Mail order gene fragments to assayable proteins within 24 hours. Biotechnology and Bioengineering 116, 667–676 (2019).

50. Parvathi, K., Chen, F., Guo, D., Blount, J. W. & Dixon, R. A. Substrate preferences of O-methyltransferases in alfalfa suggest new pathways for 3-O-methylation of monolignols. Plant Journal 25, 193–202 (2001).

51. Osakabe, K. et al. Coniferyl aldehyde 5-hydroxylation and methylation direct syringyl lignin biosynthesis in angiosperms. Proceedings of the National Academy of Sciences 96, 8955–8960 (1999).

52. Li, L., Popko, J. L., Umezawa, T. & Chiang, V. L. 5-Hydroxyconiferyl aldehyde modulates enzymatic methylation for syringyl monolignol formation, a new view of monolignol biosynthesis in angiosperms. Journal of Biological Chemistry 275, 6537–6545 (2000).

